# Placental transcriptomics reveals alterations in metabolic and transport gene programs following prenatal cannabis smoke exposure

**DOI:** 10.64898/2026.08.31.748443

**Authors:** Cristina Monaco, Tina Podinic, Donald Xhuti, Jenny S. Feeney, Doris Tan, Melanie Lemaire, Andie MacAndrew, Maria Sunil, Joshua P. Nederveen, Samantha L. Wilson, Elyanne Ratcliffe, Sandeep Raha

## Abstract

*Background:* Prenatal cannabis use is increasing, yet the effects of inhaled cannabis smoke on placental molecular function remain poorly defined. *Objective:* To characterize placental transcriptomic alterations following prenatal cannabis smoke exposure using a physiologically relevant murine inhalation model. Methods: Timed-pregnant CD-1 mice were exposed to cannabis cigarette smoke or room air from embryonic day (E)6.5 to 18.5. Placentae were collected at E18.5 for microarray-based transcriptomic profiling. Differential gene expression was assessed using fetal sex adjusted linear models and false discovery rate correction. Gene set variation and ranked pathway enrichment were performed to evaluate compartment-associated and biological transcriptional programs. Selected genes were validated by quantitative PCR. Results: Large-scale transcriptomic profiles revealed no clear visual clustering by exposure; however, 23 genes were differentially expressed (FDR <0.05, |log2 fold change| > 0.5). Labyrinth-associated transcriptional signatures were reduced, whereas junctional-associated signatures were increased in exposed placentae. Pathway analyses revealed negative enrichment of fatty acid metabolism, bioenergetic, and transmembrane transport pathways, alongside positive enrichment of RNA processing and transcriptional-regulatory pathways. Quantitative PCR confirmed altered expression of multiple transport-related genes. Electron transport chain-associated gene expression was altered despite preserved placental respiratory measurements. *Conclusions:* Prenatal cannabis smoke exposure is associated with subtle yet coordinated placental transcriptional alterations involving metabolic, transport-associated, and gene-regulatory pathways, alongside compartment-associated transcriptional shifts. These findings provide insight into placental responses to whole cannabis smoke exposure and identify pathways that may contribute to altered placental function during pregnancy.

**Graphical Abstract:** 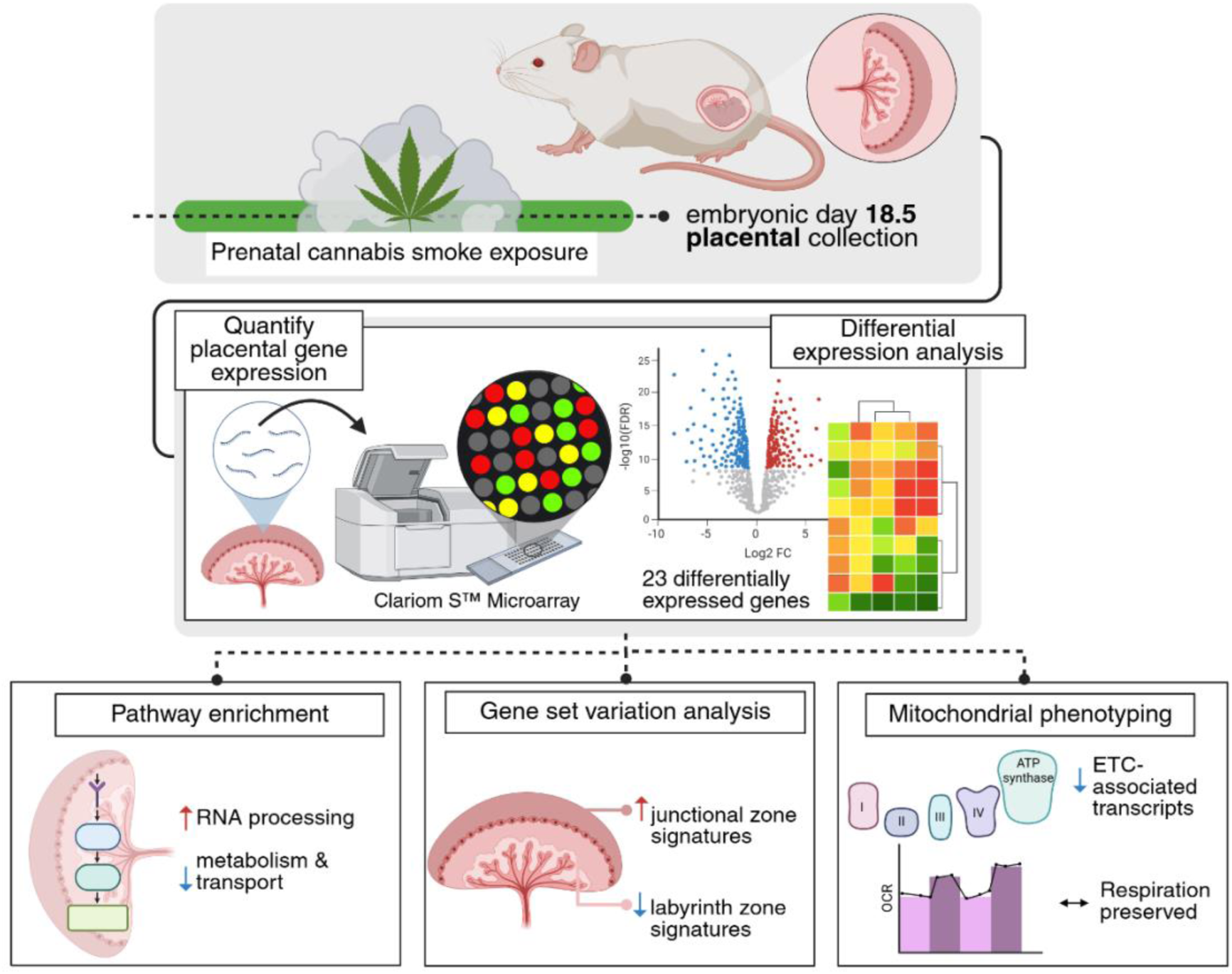

## Introduction

Cannabis is among the most commonly used psychoactive substances during pregnancy, with prenatal use continuing to rise across North America (1, 2). Importantly, prenatal cannabis use has been associated with adverse pregnancy outcomes, including fetal growth restriction and preterm birth, and smoking remains a frequently reported mode of consumption (3, 4). Despite this, many preclinical studies focus on isolated cannabinoid constituents (e.g., Δ9- tetrahydrocannabinol, Δ9-THC) administered by intraperitoneal injection, which may not accurately represent real-world exposure to cannabis smoke (5). Notably, studies comparing inhalation- and injection-based exposure models have reported differences in cannabinoid distribution in maternal circulation and fetal tissues, as well as distinct effects on maternal and litter health and offspring behavioural outcomes, underscoring the importance of exposure route (6–8). Collectively, these findings suggest that conventional preclinical models may not fully capture the biological effects of real-world cannabis use during pregnancy. Consequently, the biological mechanisms linking prenatal cannabis smoke exposure to adverse pregnancy outcomes remain poorly defined, particularly with respect to placental function.

The placenta is a highly specialized organ at the maternal-fetal interface that regulates nutrient exchange, endocrine signaling, and fetal development (9, 10). As such, it is directly exposed to circulating xenobiotics and acts as a key mediator of prenatal environmental exposures (11, 12). Accordingly, disruption of placental function may have important consequences for fetal development and pregnancy outcomes (10). Mitochondria are central regulators of placental energy metabolism and support the energy-demanding processes including nutrient transport, endocrine activity, redox homeostasis, and trophoblast differentiation. Emerging evidence from our group and others indicates that cannabis exposure can alter trophoblast mitochondrial and metabolic function (13–17). Therefore, cannabis-induced changes in mitochondrial function may contribute to impaired placental efficiency and adverse pregnancy outcomes.

Preclinical studies have shown that individual cannabinoid constituents, including Δ9-THC and cannabidiol (CBD), can disrupt trophoblast function in vitro and impair placental development in vivo, with downstream effects on nutrient transport, vascular function and fetal growth (13, 14, 18, 19). However, these approaches may not capture the complexity of whole cannabis smoke exposure, which contains a diverse mixture of cannabinoids, other bioactive compounds, and combustion-derived compounds (20). Consistent with this, emerging evidence from an in vitro cannabis smoke extract (CaSE) model demonstrates distinct effects on xenobiotic metabolism and trophoblast function, which differ from those observed with isolated cannabinoids and align with findings from in vivo inhalation-based exposure studies (8, 15).

Building on these observations, we previously reported that prenatal cannabis smoke exposure in a murine model was associated with altered placental development, including reduced placental size, altered zonation, and reduced expression of markers associated with trophoblast differentiation, endocrine function, and nutrient transport (21). These findings indicate that cannabis smoke exposure is associated with disruption of key placental processes, however, the underlying molecular mechanisms remain poorly defined. Transcriptomic profiling provides an unbiased approach to identify gene-expression changes and molecular pathways associated with placental responses to environmental exposures (22, 23). To our knowledge, no prior studies have performed genome-wide transcriptomic profiling of placental tissues following prenatal cannabis smoke exposure. Therefore, in the present study, we used microarray-based transcriptomic profiling of placental tissue from our murine inhalation model to characterize exposure-associated gene-expression changes and identify molecular pathways that may contribute to cannabis smoke-associated placental dysfunction. Given the importance of mitochondrial metabolism in placental function, we additionally assessed placental respiratory activity, electron transport chain protein content, and expression of mitochondrial respiratory genes.

### Methods Animal model

All procedures were approved by the McMaster University Animal Research Ethics Board (AUP #21-06-16). Timed-pregnant female CD-1 mice were exposed to cannabis cigarette smoke using a previously described inhalation paradigm (21, 24). Briefly, dams were exposed daily to mainstream cannabis smoke generated from the combustion of six THC-dominant cannabis cigarettes (12-14% THC; 0-2% CBD) or room air from embryonic day (E)6.5 to E18.5 using a whole-body inhalation system for 30 minutes per day. At E18.5, dams were euthanized via carbon dioxide (CO_2_) asphyxiation, and placentas corresponding to individual fetuses were collected, weighed, and snap-frozen for downstream analyses. Paired maternal and fetal liver samples were also collected to confirm fetal cannabinoid exposure by targeted quantification of cannabinoids, as previously described (21). One placenta per dam was randomly selected for downstream analyses to ensure that dam/litter was treated as the experimental unit.

### Genotyping for sex determination

Fetal sex was determined by PCR amplification of the Y-linked sex-determining region (*Sry* locus; indicative of male fetuses) and an autosomal control gene (*Fabp2).* Genomic DNA (gDNA) was extracted from snap-frozen fetal tail tissue using the Lyse-N-Amp Mouse Genotyping Kit (GeneBio Systems) according to the manufacturer’s instructions. Quantitative PCR (qPCR) was performed using the following cycling conditions: 95 °C for 10 minutes (initial denaturation), followed by 35 cycles of 95 °C for 15 sec (denaturation), 60 °C for 1:00 (annealing/extension). Dissociation melt curves were generated from 65 °C to 95°C in 1 °C increments to confirm amplification specificity. Primer sequences are provided in Supplementary Table S1.

### RNA Extraction & processing

Snap-frozen E18.5 placentae (∼50mg) were mechanically homogenized in 1 mL of ice- cold TRIzol^TM^ reagent (Thermo Fisher Scientific). Total RNA was then isolated using the Direct- zol RNA Extraction Kit (Zymo Research, R2060) according to the manufacturer’s instructions. RNA concentrations (ng/μl) and purity were assessed using a Implen NanoPhotometer N60. RNA integrity was evaluated using a Bioanalyzer with the RNA 6000 Nano Kit (Agilent Technologies). Samples with an RNA integrity (RIN) ≥ 8 were used for downstream microarray analysis.

### Microarray processing & normalization

Microarray profiling was performed using the Affymetrix Clariom^TM^ S Mouse Array (Thermo Fisher Scientific), which interrogates >21,000 well-annotated mouse genes. Briefly, 200 ng of total RNA per sample was processed for labeling, hybridization, and scanning according to the manufacturer’s standard GeneChip protocol carried out at the Population Health Research Institute CRLB-GMEL facility, McMaster University. Raw CEL files were generated and imported into Transcriptome Analysis Console (TAC) software version 4.0.3.14 for quality control assessment and preprocessing. Gene expression data were normalized using the Signal Space Transformation-Robust Multi-Array Average (SST-RMA) algorithm which performs background correction, signal-space transformation, quantile normalization, and probe set summarization to generate log2-transformed gene expression values for downstream analysis (25, 26). Gene-level features were considered expressed if their detection above background (DABG) values were <0.05 in at least 50% of samples. The manufacturer-provided annotation file, *Clariom_S_Mouse.r1.na36.mm10.a1.transcript.csv*, based on the mm10 mouse reference genome, was used for gene-level feature annotation.

### Differential expression analysis & probe annotation

R version 4.5.2 (R Core Team, 2024) was used for all downstream data analyses (27). Normalized log2 gene expression values between cannabis exposed (n = 7) and control (n = 7) placental samples were compared with a linear regression model adjusting for fetal sex in limma (version 3.66.0) (25).

Moderated t-statistics were calculated using empirical Bayes variance estimation (28). The p-values were adjusted for multiple testing using the Benjamini-Hochberg false discovery rate (FDR) method. Genes were considered differentially expressed if they had an FDR <0.05 and |log2fold change| > 0.5. Of the 22,206 probes/features tested, 20,995 were annotated with gene symbols. In instances where multiple probes mapped to the same gene, expression values were collapsed to a single gene-level value by retaining the probe with the highest average expression across samples, resulting in 20,737 unique gene-level entries for downstream visualization and pathway analysis.

### Data visualization (PCA, volcano, heatmaps)

Principal component analysis (PCA) was conducted on normalized log2-transformed gene-level expression values, centered and scaled across samples, to assess overall transcriptional variation. Associations between principal component scores and exposure group or fetal sex were assessed using linear models including both variables as predictors. Heatmaps of significantly differentially expressed genes (FDR < 0.05 and |log2 fold change| > 0.5) were generated using normalized log2-transformed expression values, row-scaled to z-scores to visualize relative expression across samples using pheatmap (version 1.0.13) (29). Samples and genes were hierarchically clustered using correlation distance and complete linkage.

### Gene set variation analysis (GSVA)

Gene set variation analysis (GSVA) was performed to estimate sample-wise enrichment scores for curated gene sets representing placental compartment-associated transcriptional programs using GSVA (version 2.4.7) (30). Gene sets were manually curated from published single-cell and single-nuclei RNA sequencing studies of the mouse placenta listed in **Supplementary Table S3** (31–34). GSVA scores were computed using a non-parametric kernel estimation method with a Gaussian distribution assumption and default parameters. Differences in GSVA enrichment scores between control and cannabis smoke-exposed groups were assessed using linear models adjusted for fetal sex.

### Gene set enrichment and pathway analysis

Gene set enrichment analysis (GSEA) was performed on the 20,737 unique Entrez-mapped genes using clusterProfiler (version 4.18.4) (35, 36). Genes were ranked according to the moderated t-statistic obtained from the limma model, preserving both the direction and magnitude of differential expression. Gene sets were evaluated using only genes represented in the retained, Entrez-mapped ranked microarray gene list. Thus, enrichment was assessed relative to the array-derived expressed gene universe rather than the entire mouse genome. This rank-based approach enables the detection of pathway-level changes across the full gene-expression distribution without applying an arbitrary differential-expression significance threshold (37).

Enrichment analysis was conducted using gene sets from the following databases: Gene Ontology (GO) biological processes, Kyoto Encyclopedia of Genes and Genomes (KEGG), Reactome pathways, and the Molecular Signatures Database (MSigDB), including the hallmark (H) collection (38–41). We included all databases as they provide complementary biological annotations at different levels of granularity. Results across databases were interpreted as complementary annotations of shared biological themes rather than as independent evidence of pathway alteration. Redundant terms were reduced using similarity-based simplification, with highly similar terms removed at a similarity cutoff of 0.7 while retaining the term with the lowest FDR. Gene sets with an FDR < 0.05 were considered significantly enriched.

### Quantitative PCR

Total RNA (1 μg) was reverse-transcribed into cDNA using the Applied Biosystems^TM^ High-Capacity cDNA Reverse Transcription Kit (Applied Biosystems, 4368813) according to the manufacturer’s protocol. Quantitative PCR (qPCR) was performed using the TouchTM Real-Time PCR Detection System with GB-Amp™ InFluor^TM^ qPCR reagent on a Bio-Rad CFX384 platform (GeneBio, P2092). Relative gene expression was calculated using the ΔΔCt method and normalized to Actb as the housekeeping gene (42). Primer sequences used in the qPCR validation are listed in **Supplementary Table S1.**

### Mitochondrial respirometry from placental tissue

Complex-associated respiration was measured in frozen mouse placental tissues using a previously established protocol (43). Briefly, ∼100 mg of mouse placental tissue was mechanically homogenized in ice-cold 1X mitochondrial assay (1X MAS) buffer. Mouse placental homogenates were quantified for total protein using the Pierce^TM^ BCA Protein Assay kit (Thermo Fisher, #23225) according to the manufacturer’s protocol. Placental homogenates were further processed via differential centrifugation to obtain placental mitochondrial-enriched fractions (MEFs). Following normalization to whole placental homogenate protein abundance, 100 µl of placental MEFs from CTRL and EXP groups (n = 10/group) were loaded into duplicate wells of a Seahorse XF24 microplate and suspended in additional 1X MAS buffer containing a cocktail of respiratory substrates (1mM NADH, 4mM ADP, 10mM pyruvate, 1mM malate). Exogenous NADH was included to support Complex I activity in the membrane-disrupted preparation, whereas pyruvate and male were supplied to support endogenous NADH generation in mitochondria with residual membrane integrity and to maintain consistency with the established assay protocol.Fi To measure complex I-, II- and IV-associated oxygen consumption rates (OCR) in a single protocol, each of four ports in the pre-hydrated cartridge was loaded (50µl/port) with a sequence of substrate-inhibitor combinations as follows (ports A: rotenone; B: succinate; C: antimycin A; D: TMPD/ascorbate). The mix/wait/measure times were set to 0.5 min/0.5 min/2 min under the “Complex-Associated” protocol. All respirometry experiments were conducted using biologically independent placental samples from separate dams. OCR readouts from duplicate wells were averaged per biological replicate and represented in pmol/min/µg.

### Immunoblotting

Placental MEFs corresponding to respirometry experiments were sonicated on ice for three rounds of five pulses at 5 Hz. Protein lysates were quantified for protein using the Pierce^TM^ BCA Protein Assay kit (Thermo Fisher, #23225) according to the manufacturer’s protocol. Protein lysates were prepared at a final concentration of 1mg/ml using 6X sample buffer. Mitochondrial fractions were loaded (10µg/well) and resolved on a 4-20% Criterion TGX^TM^ Stain-Free^TM^ Protein Gel (BioRad). Protein was subsequently transferred to LF-PVDF membranes using the semi-dry Trans-Blot Turbo System (BioRad). Membranes were blocked for 1 hour at RT and incubated in primary antibodies at 4°C overnight in 5% bovine serum albumin (BSA) in 1X TBST solutions: Total OXPHOS Rodent Antibody Cocktail (1:1000 dilution; Abcam, ab110413); anti-CS antibody (1:1000 dilution; custom-made). 1:10000 dilutions of either anti-mouse or rabbit horseradish peroxidase (HRP)-linked secondary antibodies were incubated for 1 hour at RT in 5% BSA in 1X TBST. Membranes were briefly incubated with Bio- Rad Clarity Max Western ECL Substrate and imaged on the ChemiDoc Imaging System (BioRad). Densitometric analyses were performed using the ImageLab software (BioRad) and normalized to stain-free images of total protein.

### Statistical Analyses

All statistical analyses were performed in R (version 4.5.2) and GraphPad Prism version 10.4.0. For transcriptomic analyses, differential expression, gene set variation analysis, and pathway enrichment were performed in R as described above. For transcriptomic analyses, statistical significance was determined using the Benjamini-Hochberg false discovery rate method. For qPCR validation and selected two-group comparisons, differences between control and cannabis smoke-exposed groups were assessed using Welch’s unpaired two-tailed t-test. For respiratory measurements, data were analyzed using one-way ANOVA, followed by a post-hoc Tukey test. Data are presented as mean ± standard error of the mean (SEM) unless otherwise indicated, and statistical significance was defined as p <0.05.

## Results

### Whole-placenta transcriptomes show no pronounced large-scale separation by exposure at E18.5

Following quality control and normalization, principal component analysis (PCA) was performed to visualize large-scale patterns in whole-placenta transcriptomes. The first two principal components explained 14.2% and 10.5% of the total variance, respectively (**Figure 1A**). Samples did not exhibit clear visual separation by cannabis smoke exposure or fetal sex. Following SST-RMA preprocessing, normalized log2-transformed expression values showed comparable distributions across all samples, indicating successful between-array normalization (**Figure 1B**).

**Figure 1.**
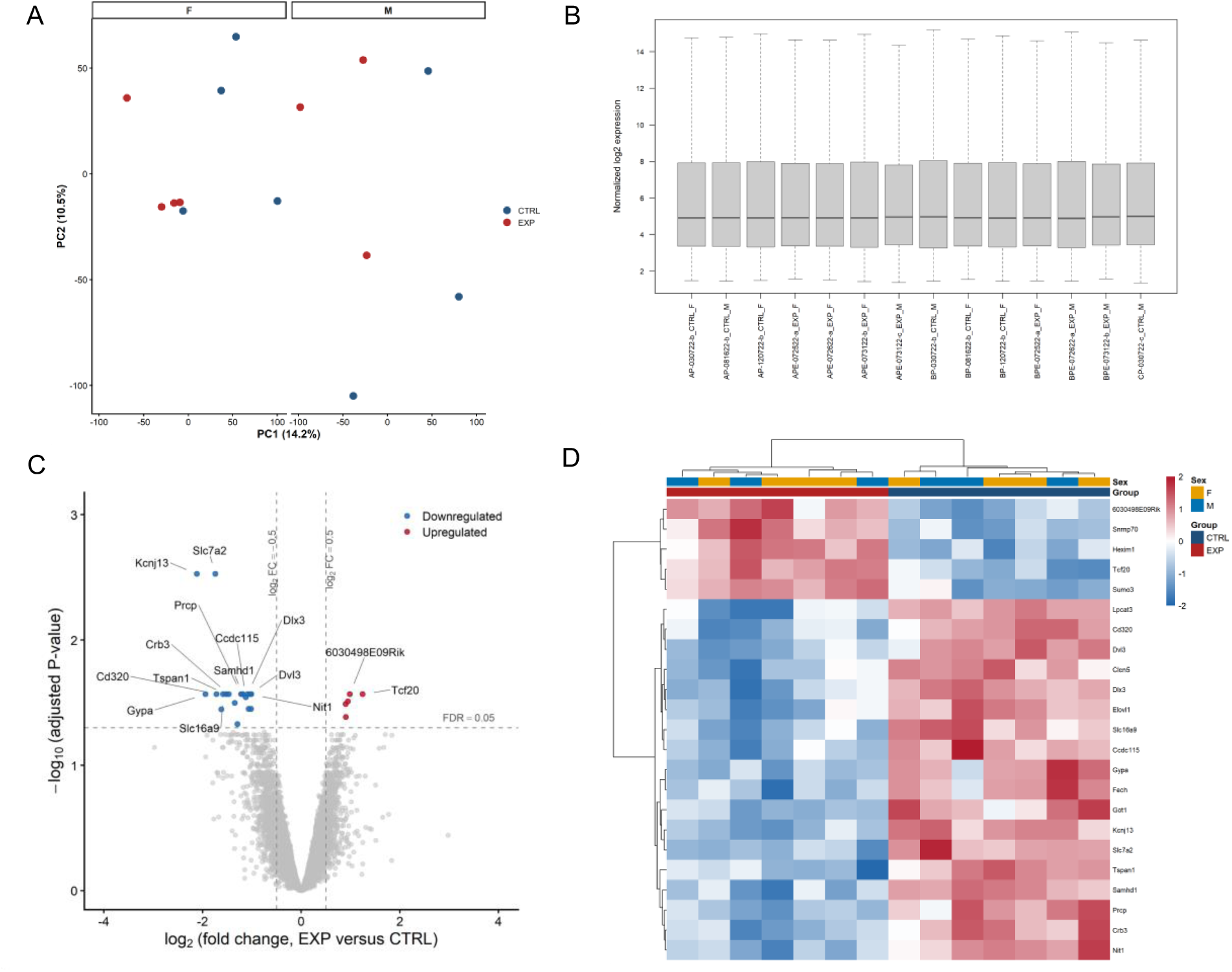
Prenatal cannabis smoke exposure induces subtle placental transcriptomic changes involving transport- and regulatory-associated genes. **(A)** Principal component analysis (PCA) of normalized log2-transformed placental transcriptomes from control (CTRL) and cannabis smoke-exposed (EXP) placentae at E18.5. Samples are coloured by exposure group and separated by fetal sex. PC1 and PC2 explained 14.2% and 10.5% of the total variance, respectively. **(B)** Distribution of normalized log2 expression values across individual placental samples following SST-RMA preprocessing. **(C)** Volcano plot of differential expression in E18.5 placentae from CTRL and EXP groups. Differential expression was tested across 22,206 retained microarray probes/features using limma with fetal sex included as a covariate. Following annotation and gene-level collapsing, 20,737 gene-level features were retained for visualization and downstream interpretation. Dashed vertical lines indicate |log2 fold change| = 0.5, and the dashed horizontal line indicates FDR-adjusted p = 0.05. Representative significantly dysregulated genes are annotated. **(D)** Heatmap of significantly differentially expressed genes in E18.5 placentae from CTRL and EXP groups. Normalized log2 expression values were adjusted for fetal sex while preserving the cannabis smoke exposure-group effect, then row-scaled by gene to z-scores. Columns represent individual placental samples and are annotated by fetal sex and exposure group. Rows represent significantly differentially expressed genes meeting FDR- adjusted p < 0.05 and |log2 fold change| > 0.5.

Unsupervised hierarchical clustering of the 1,000 genes with the greatest variance in normalized log2-transformed expression values did not reveal distinct segregation by exposure group or fetal sex (**Figure S1**), supporting the absence of pronounced large-scale separation of whole-placenta transcriptomes by these variables.

### Differential gene-expression analysis identifies a subset of placental genes altered by prenatal cannabis smoke exposure

Despite the absence of large-scale transcriptomic separation, we next sought to identify specific genes altered by cannabis smoke exposure. We assessed differential gene-expression patterns between cannabis smoke-exposed and control placentae, with fetal sex included as a covariate. Statistically significant differentially expressed genes were defined as having an FDR <0.05 and |log2fold change| > 0.5. We identified 23 significantly differentially expressed genes between exposed (EXP) and control (CTRL) placentae, of which 19 genes were downregulated and 4 were upregulated in the exposed group (**Figure 1C; Supplementary Table S2**). Volcano plot visualization indicated that most significantly differentially expressed genes exhibited modest fold changes, consistent with the absence of large-scale transcriptomic shifts (**Figure 1C**). Among the most downregulated genes included *Kcnj13*, *Slc7a2*, and *Gypa*, while *Tcf20* and *Hexim1* were among the few genes upregulated following exposure. These genes are associated with ion transport, amino-acid transport, erythroid-related pathways, and transcriptional regulation, consistent with the pathway-level alterations identified in subsequent analyses. Hierarchical clustering revealed distinct expression patterns between cannabis smoke-exposed and control placentae, with CTRL and EXP samples clustering separately despite within-group variation (**Figure 1D**).

### Cannabis smoke exposure alters placental compartment-associated transcriptional signatures

Given the central roles of the labyrinth and junctional zones in maternal-fetal exchange, placental endocrine function, and trophoblast differentiation, we performed a placental regional compartment-focused analysis using curated gene signatures derived from murine placental single-cell RNA-sequencing datasets. Two curated gene sets, comprising a total of 15 genes, were used to represent labyrinth- and junctional zone-associated transcriptional programs (**Supplementary Table 3)**. These gene sets included established cell-identity markers and transcriptionally associated genes for the labyrinth and junctional zones, the two major functionally distinct compartments of the mouse placenta. The labyrinth zone mediates maternal-fetal gas, nutrient, and waste exchange, whereas the junctional zone is primarily involved in trophoblast differentiation, endocrine signaling, and structural support. Gene set variation analysis generated sample-level enrichment scores for each compartment-associated gene set.

After adjustment for fetal sex, labyrinth zone-associated gene signatures were significantly reduced in cannabis smoke-exposed placentae compared with controls (**Figure 2A**; adjusted p < 0.0001). In contrast, junctional zone-associated signatures were significantly increased following exposure (**Figure 2B**; adjusted p = 0.0404). These findings indicate exposure-associated differences in labyrinth- and junctional zone-associated transcriptional programs. As GSVA scores reflect coordinated gene-expression patterns, they should not be interpreted as direct measures of placental zone size, cell abundance, or tissue composition.

**Figure 2.**
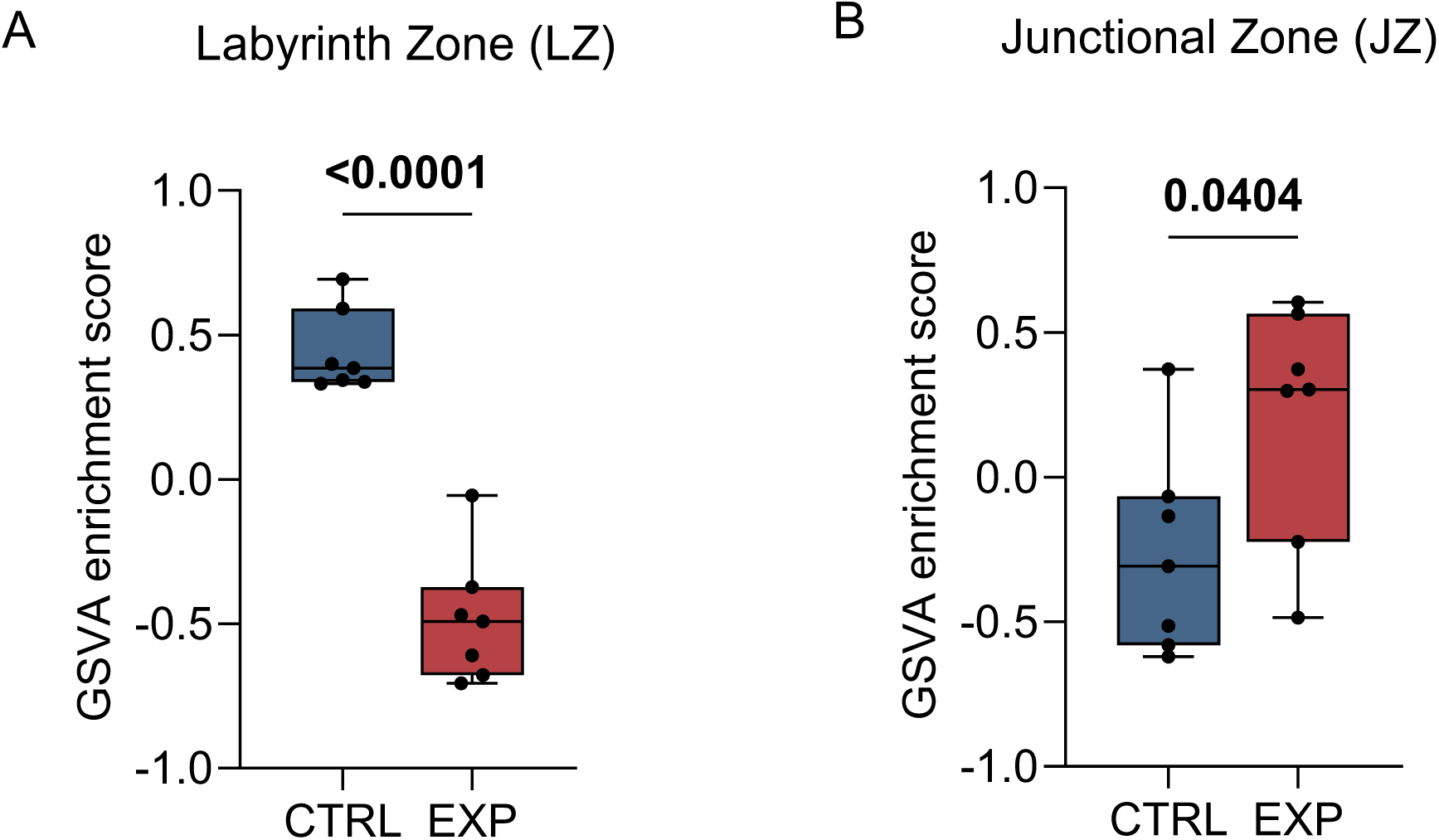
Prenatal cannabis smoke exposure is associated with divergent placental regional-associated transcriptional signatures. **(A)** Labyrinth zone (LZ)-associated transcriptional signatures were reduced in cannabis smoke-exposed (EXP) placentae compared with controls (CTRL). **(B)** Junctional zone (JZ)-associated transcriptional signatures were increased following exposure. Gene set variation analysis (GSVA) was performed on bulk placental microarray data using curated marker gene sets representing labyrinth- and junctional zone-associated transcriptional programs. Higher GSVA enrichment scores indicate stronger relative coordinated expression of genes within the corresponding compartment-associated signature, whereas lower scores indicate reduced relative expression of that transcriptional program. Enrichment scores are shown for CTRL and EXP groups as boxplots with individual samples overlaid. Differences in GSVA enrichment scores were assessed using limma linear models with exposure group as the main predictor and fetal sex included as a covariate. P values were adjusted across the two tested signatures using the Benjamini–Hochberg false discovery rate method.

### Pathway enrichment analysis reveals coordinated alterations in metabolic, transport-associated and gene-regulatory pathways following prenatal cannabis smoke exposure

To determine whether biological pathways were altered despite modest gene-level changes, gene set enrichment analysis (GSEA) was performed using a ranked gene list on the moderated t-statistic derived from differential expression analysis. Gene Ontology (GO) biological process analysis revealed positive enrichment of pathways related to regulation of mRNA processing in cannabis smoke-exposed placentae. In contrast, pathways involved in lipid modification, cholesterol metabolism, membrane lipid biosynthesis, organic cation transport and transmembrane transport were negatively enriched (**Figure 3A).** Consistent with these findings, KEGG pathway analysis identified positive enrichment of the spliceosome pathway, alongside negative enrichment of pathways associated with carbon metabolism and fatty acid metabolism in exposed placentae (**Figure 3B).** Reactome pathway analysis further supported these patterns, demonstrating negative enrichment of pathways related to lipid metabolism, solute carrier-mediated transport, aerobic respiration, respiratory electron transport, and oxidative phosphorylation-related processes in exposed samples (**Figure 3C).** Hallmark gene set enrichment analysis revealed negative enrichment of signaling pathways related to oxidative phosphorylation, fatty acid metabolism, adipogenesis, xenobiotic metabolism, IL-2-STAT5 signaling, and mTORC1 signaling (**Figure 3D).** Together, these findings indicate alterations in metabolic, bioenergetic, transport-associated and gene-regulatory transcriptional programs following prenatal cannabis smoke exposure.

**Figure 3.**
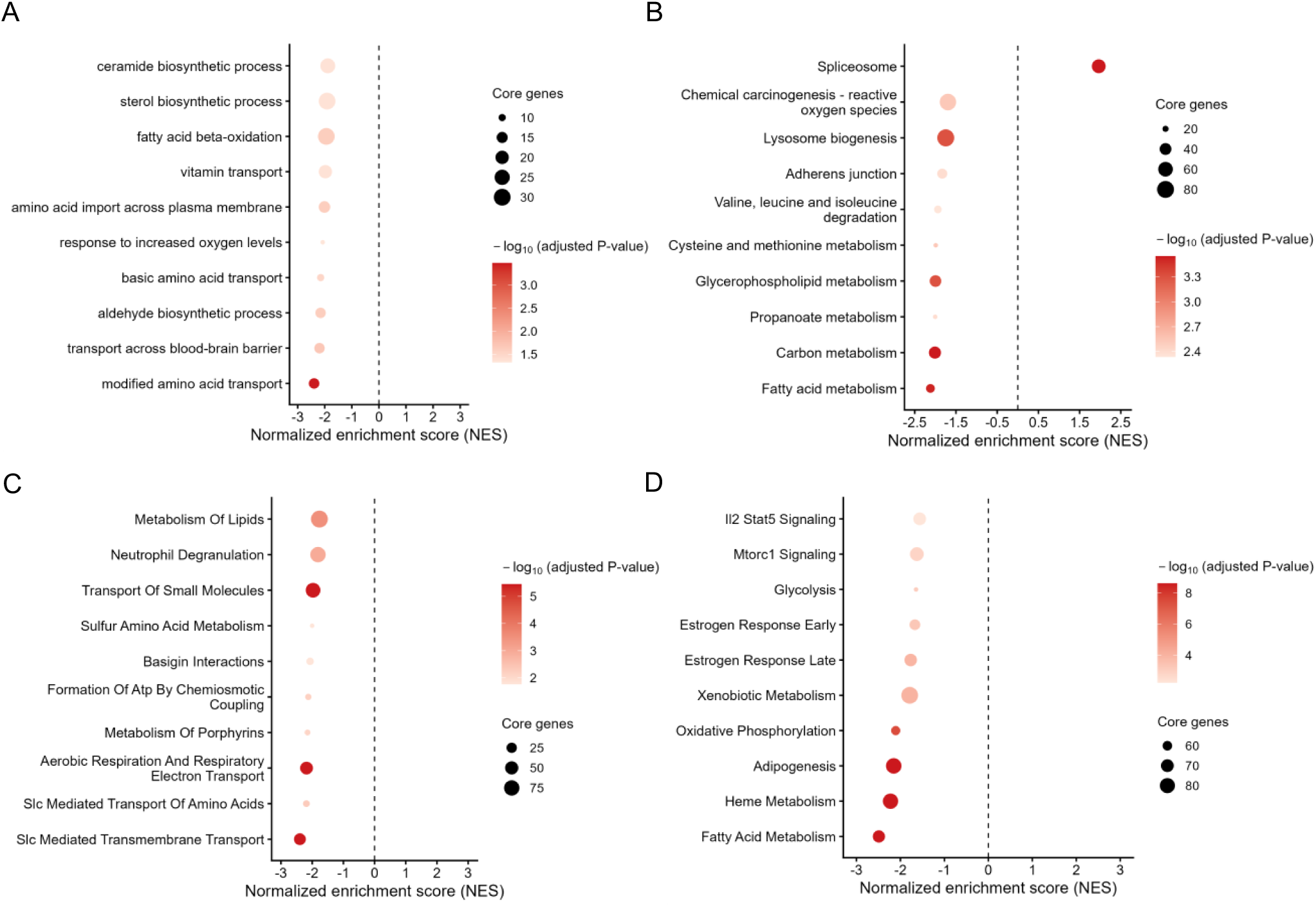
Prenatal cannabis smoke exposure is associated with coordinated enrichment of RNA-processing pathways and negative enrichment of metabolic, bioenergetic, and transport pathways. **(A)** Gene Ontology (GO) biological process enrichment analysis. **(B)** Kyoto Encyclopedia of Genes and Genomes (KEGG) pathway enrichment analysis. **(C)** Reactome pathway enrichment analysis. **(D)** Hallmark gene set enrichment analysis. Enrichment analyses were performed using ranked gene lists based on the moderated t-statistic from differential-expression analysis. The normalized enrichment score (NES) reflects the direction and magnitude of enrichment. Positive NES values indicate enrichment among genes with higher expression in cannabis smoke-exposed placentae relative to controls, whereas negative NES values indicate enrichment among genes with lower expression in cannabis smoke-exposed placentae. Dot size corresponds to the number of core enriched genes, and colour intensity reflects statistical significance [-log10(FDR-adjusted p value)].

### qPCR validation confirms exposure-associated changes in selected differentially expressed genes

To validate differential expression identified by microarray analysis, qPCR was performed for genes selected based on their statistical significance, effect size, and biological relevance to placental nutrient transport and cellular function. The genes selected for validation were *Kcnj13, Slc7a2, Slc16a9, Gypa, Tcf20* and *Clcn5*. qPCR validation was performed in an expanded cohort that included the placental RNA samples used for microarray analysis, comprising control placentae (CTRL; n = 15; 7 male and 8 female) and cannabis smoke-exposed placentae (EXP; n = 10; 5 male and 5 female). Consistent with transcriptomic findings, several transport-related genes were significantly reduced in cannabis smoke-exposed (EXP) placentae compared with controls (CTRL), including *Clcn5* (p=0.0168), *Kcnj13* (p=0.0090), *Gypa* (p=0.0029), *Slc7a2* (p=0.0005), *Slc16a9* (p=0.0285), (**Figure 4A-E).** In contrast, *Tcf20* expression was significantly increased following exposure (**Figure 4F**; p=0.0059). These findings support the directionality of microarray-derived differential-expression results.

**Figure 4.**
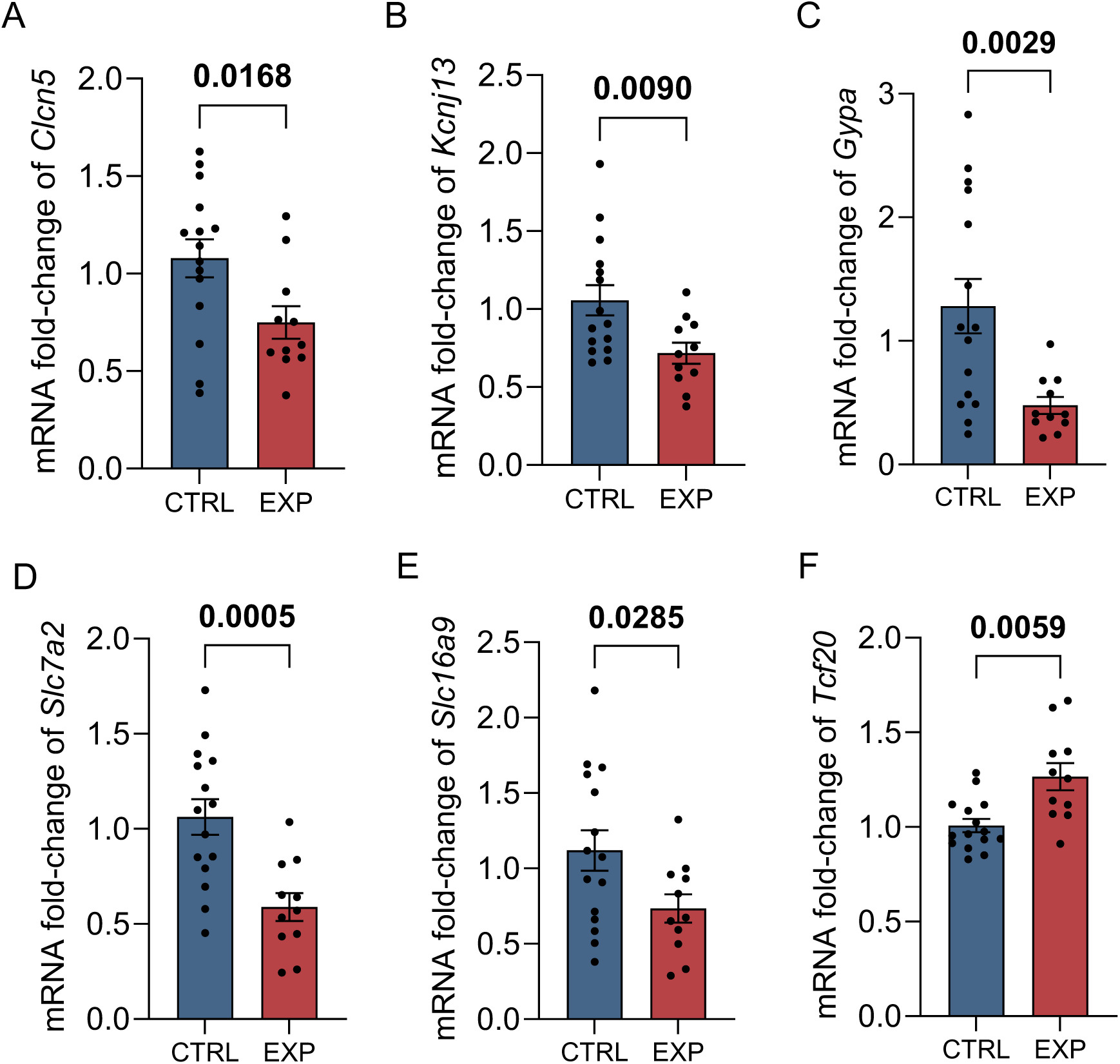
qPCR validation confirms exposure-associated changes in selected transport- and regulatory-associated genes. Relative expression of (A) *Clcn5*, (B) *Kcnj13*, (C) *Gypa*, (D) *Slc7a2*, (E) *Slc16a9*, and (F) *Tcf20* in control (CTRL; n = 15, 7 male and 8 female) and cannabis smoke-exposed (EXP; n = 10, 5 male and 5 female) placentae. Gene expression was normalized to *Actb*. Data are presented as mean ± SEM with individual samples overlaid. Statistical significance was determined using Welch’s unpaired two-tailed t-test, with p values displayed.

### Prenatal cannabis exposure alters placental mitochondrial gene expression without changes in respiratory capacity or oxidative phosphorylation protein content

Given the negative enrichment of bioenergetic, respiratory electron transport, and oxidative phosphorylation-related pathways identified by pathway analysis, we next examined whether prenatal cannabis smoke exposure was associated with altered placental mitochondrial gene regulation and respiratory function. We first assessed the expression of mitochondrial respiratory-chain genes in an expanded cohort that included the placental RNA samples used for microarray analysis, comprising control placentae (CTRL; n = 15; 7 male and 8 female) and cannabis smoke-exposed placentae (EXP; n = 10; 5 male and 5 female). Several electron transport chain-associated transcripts were altered in cannabis smoke-exposed placentae compared with controls. Specifically, *Ndufb8* was significantly reduced in exposed placentae (p = 0.0172), while *Cox4i1* (p = 0.0274), *Cox5a* (p = 0.0169), and *Atp5f1a* (p = 0.0169) were also significantly decreased (Figure 5A). In contrast, *Sdhb* was significantly increased following exposure (p = 0.0473), and *Ndufv1* also appeared increased (**Figure 5A**). Interestingly, not all respiratory enzyme complex-associated subunits demonstrated an increase in mRNA expression, suggesting selective changes in mitochondrial gene regulation rather than uniform alterations of electron transport chain gene expression **(Figure 5A)**. To determine whether these transcript-level changes were associated with altered mitochondrial respiratory capacity, complex-associated oxygen consumption rates were measured in mitochondrial-enriched placental fractions (MEFs) isolated from a separate set of frozen E18.5 placentae from cannabis smoke-exposed (EXP; n=10, 5 female and 5 male) and control (CTRL; n=10, 5 female and 5 male) groups. Representative oxygen-consumption traces showed the expected responses to sequential substrate and inhibitor additions used to assess complex I-, II-, and IV-associated respiration (**Figure 5B**). Quantification of complex-associated respiration revealed no significant differences between control and cannabis smoke-exposed placentae (**Figure 5C**). We next assessed whether prenatal cannabis smoke exposure altered electron transport chain protein abundance.

**Figure 5.**
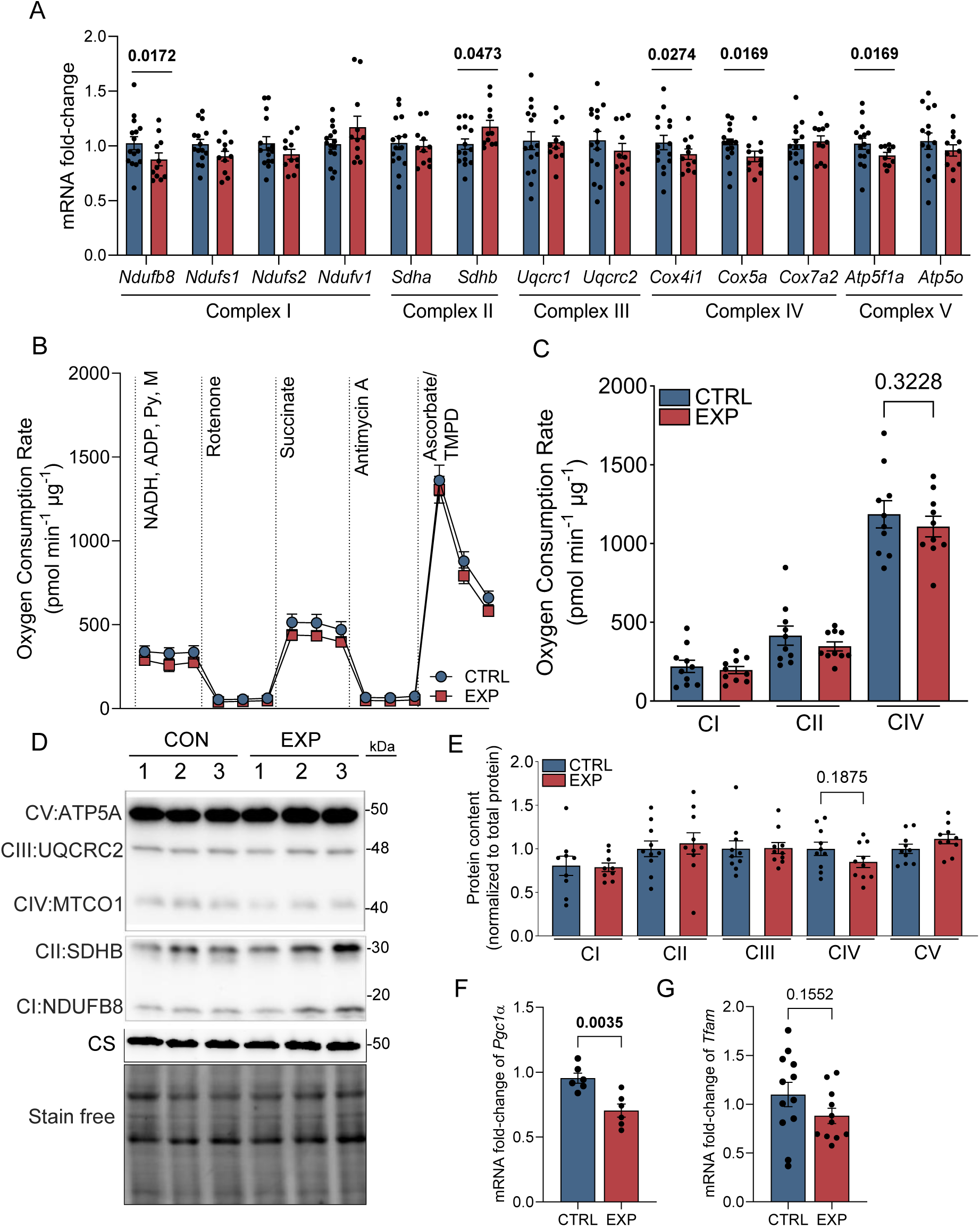
**Prenatal cannabis smoke exposure alters placental mitochondrial gene regulation without detectable changes in respiratory capacity or electron transport chain protein content**. **(A)** Relative mRNA expression of mitochondrial respiratory-chain subunits in control (CTRL) and cannabis smoke-exposed (EXP) whole placental lysates, normalized to *Actb*. Genes are grouped by electron transport chain complex. **(B)** Representative oxygen-consumption-rate (OCR) traces measured in mitochondrial-enriched fractions (MEFs) isolated from E18.5 mouse placentae following prenatal exposure to room air (CTRL) or cannabis smoke (EXP). Sequential substrate and inhibitor injections were used to assess complex I (CI)-, complex II (CII)-, and complex IV (CIV)-associated respiration. **(C)** Quantification of CI-, CII-, and CIV-associated OCR in CTRL and EXP placental MEFs. **(D)** Representative immunoblots of oxidative phosphorylation (OXPHOS) complex subunits, including CI: NDUFB8, CII: SDHB, CIII: UQCRC2, CIV: MTCO1, CV: ATP5A, and citrate synthase (CS), with stain-free total protein shown as a loading control. **(E)** Quantification of OXPHOS subunit protein content in CTRL and EXP placental MEFs, normalized to stain-free total protein. (F, G) Relative mRNA expression of (F) *Ppargc1a*, encoding PGC-1α, and (G) *Tfam* in CTRL and EXP whole placental lysates, normalized to *Actb*. Data are presented as mean ± SEM with individual samples overlaid. Statistical significance was determined using Welch’s unpaired two-tailed t-test for two-group comparisons or one-way ANOVA with Tukey’s post hoc test for respiratory measurements. p < 0.05 was considered statistically significant. Py, pyruvate; M, malate; TMPD, N,N,N′,N′- tetramethyl-p-phenylenediamine.

Representative immunoblots showed detectable oxidative phosphorylation complex subunits corresponding to complexes I–V, with citrate synthase and stain-free total protein included for normalization (**Figure 5D**). Quantification revealed no significant exposure-associated differences in OXPHOS complex protein content in exposed placentae (**Figure 5E**). Finally, we examined markers of mitochondrial biogenesis and transcriptional regulation. Expression of *Ppargc1a*, encoding PGC-1α, was significantly reduced in cannabis smoke-exposed placentae (p = 0.0035; **Figure 5F**), whereas *Tfam* showed a non-significant trend toward reduced expression (p = 0.1552; **Figure 5G**). Together, these findings indicate that prenatal cannabis smoke exposure is associated with altered placental mitochondrial gene regulation, including reduced expression of selected electron transport chain-associated transcripts and *Ppargc1a*, despite preserved complex-associated respiratory capacity and OXPHOS protein content at E18.5.

## Discussion

Despite growing clinical and public health concerns surrounding cannabis use during pregnancy, the effects of prenatal cannabis smoke exposure on placental molecular function remain incompletely understood (44). Using our murine model, we previously found that cannabis smoke exposure was associated with increased feto-placental weight ratio, reduced placental weight, and zone-specific alterations in placental architecture, alongside changes in expression of markers related to placental development (21). Building on these findings, the present study used whole-placenta transcriptomic profiling to identify exposure-associated molecular changes. Although principal component analysis did not reveal pronounced separation between control and cannabis smoke-exposed placentae, differential-expression and pathway-level analyses identified subtle but coordinated transcriptional changes. These included alterations in compartment-associated signatures, metabolic and transport-related pathways, RNA-processing programs, and selected mitochondrial respiratory-chain transcripts. Importantly, these transcriptional changes occurred despite preserved complex-associated respiratory capacity and oxidative phosphorylation protein content, suggesting that prenatal cannabis smoke exposure is associated with early molecular alterations rather than overt mitochondrial respiratory dysfunction.

Given our prior observation of altered placental zonation following prenatal cannabis smoke exposure, we examined whether transcriptional programs associated with the major structural compartments of the mouse placenta differed between groups (21). Gene set variation analysis (GSVA), using gene sets representative of murine placental structural zones, revealed decreased enrichment of labyrinth zone-associated gene signatures and increased enrichment of junctional zone-associated signatures in exposed placentae **(Figure 2)**. Since the labyrinth layer is the primary site of maternal-fetal exchange and the junctional zone is more closely associated with endocrine function, these findings suggest altered compartment-associated transcriptional programs (45). Interestingly, although we have previously reported reduced expression of markers associated with murine placental differentiation and endocrine regulators, consistent with the transcriptional patterns observed in the present study, our histological analyses indicated an increased labyrinth area along with a decreased junctional zone area (21). This apparent discrepancy highlights the distinction between molecular signatures and tissue morphology.

While GSVA estimates coordinated expression of predefined marker genes in bulk tissue, it does not directly measure placental zone area, cell abundance, or cellular composition. Thus, the observed shifts here may reflect altered cellular composition or within-cell transcriptional regulation. Consistent with altered trophoblast-associated transcriptional programs, *Dlx3* was significantly downregulated in cannabis-smoke exposed placentae. Dlx3 is a placentally expressed transcription factor that holds particular importance in trophoblast differentiation and labyrinth development; whereby complete loss of Dlx3 in mice results in placental failure and abnormalities within the labyrinth compartment (46, 47). Additionally, reduced DLX3 expression is associated with decreased expression of differentiation markers, including β-hCG, syncytin, and 3β-hydroxysteroid dehydrogenase (48). Therefore, reduced Dlx3 expression in our study may reflect altered regulation of trophoblast differentiation. Nonetheless, our observations of compartment-associated shifts were detectable despite the absence of large-scale separation in the bulk transcriptomic analyses, suggesting that spatially or lineage-associated transcriptional responses may be obscured when the whole heterogeneous placenta is analyzed.

To place these compartment-associated changes within a broader molecular context, we next examined genome-wide differential-expression patterns and ranked pathway enrichment. Several transport-, nutrient-handling-, ion channel-, and iron metabolism-associated genes were downregulated in exposed placentae, whereas genes involved in transcriptional regulation and RNA processing were upregulated (**Figure 1**). Consistently, pathway-level analyses showed negative enrichment of lipid metabolic, bioenergetic, and transport-associated processes, alongside positive enrichment of RNA-processing and transcriptional-regulatory pathways (**Figure 3**). These findings suggest that prenatal cannabis smoke exposure is associated with coordinated changes in metabolic, transport-related, and gene-regulatory transcriptional programs.

The positive enrichment of pathways related to RNA processing is consistent with activation of regulatory programs under cellular stress conditions (22). Similar placental transcriptional patterns have been reported in the context of prenatal maternal stress, including enrichment of protein-processing and endoplasmic reticulum (ER) stress pathways, although whether these changes reflect adaptive or compensatory responses remains unclear (22). In the present study, gene-level analyses identified increased expression of *Hexim1* and *Snrnp70,* which are involved in transcriptional regulation and RNA splicing, respectively, as well as *Tcf20,* a transcriptional co-regulator implicated in stress-response gene expression programs (49, 50).

Previous studies have also shown that Δ9-THC and other phytocannabinoids such as CBG and CBDV can induce ER stress and activate the unfolded protein response (UPR) in trophoblast cells, a pathway known to influence transcriptional and post-transcriptional regulation (51–53). Although ER stress was not directly assessed, these findings provide a potential context for the RNA-processing and transcriptional-regulatory signatures observed following cannabis smoke exposure.

The placenta is a highly metabolically active organ that responds to environmental stress while maintaining energy-dependent functions, including nutrient transport and maternal-fetal exchange. In the present study, pathway analyses revealed negative enrichment of several metabolic and bioenergetic pathways, including oxidative phosphorylation, fatty acid metabolism, lipid metabolism, respiratory electron transport, solute transport, and membrane-associated metabolic pathways (**Figure 3**). The recurrence of related pathways across KEGG, Hallmark, Reactome, and GO analyses supports a broader pattern of altered placental metabolic and mitochondrial-associated transcriptional regulation following prenatal cannabis smoke exposure. Consistent with these enrichment patterns, targeted follow-up analysis identified altered expression of selected mitochondrial respiratory-chain transcripts, including reduced expression of *Ndufb8*, *Cox4i1*, *Cox5a*, and *Atp5f1a*, increased *Sdhb* (**Figure 5A**), and reduced *Ppargc1a* expression, with a non-significant trend toward reduced *Tfam* expression (**Figure 5F,G**). These findings suggest altered mitochondrial gene regulation following prenatal cannabis smoke exposure. However, complex-associated oxygen-consumption measurements in mitochondrial-enriched placental fractions did not reveal significant differences in respiratory capacity, and oxidative phosphorylation protein content was not significantly altered (**Figure 5B–E**). The divergence between altered mitochondrial transcript expression and preserved complex-associated respiration suggests that prenatal cannabis smoke exposure is associated with mitochondrial transcriptional remodeling in the absence of respiratory dysfunction, which is consistent with the observed shifts in bioenergetic and oxidative phosphorylation-related pathways at the transcriptomic level. Given that GSVA revealed compartment-associated transcriptional changes, it is possible that cell type- or zone-specific mitochondrial responses were not fully resolved by whole-placenta analyses. This is supported by evidence that mouse placental regions and trophoblast populations differ in energetic demands, metabolism and mitochondrial profiles, which could contribute to the absence of detectable functional differences in bulk placental measurements (54–56). Single-cell studies further demonstrate that placental cellular heterogeneity can influence bulk gene-expression profiles, supporting the need for cell-type or region-specific approaches to detect localized mitochondrial responses (57).

The enrichment patterns were also accompanied by altered expression of genes involved in lipid metabolism, membrane homeostasis, and transport. For example, *Fech* and *Lpcat3* were downregulated in exposed placentae, suggesting potential changes in heme biosynthesis and phospholipid remodeling, respectively. These processes are relevant to mitochondrial ETC function, membrane homeostasis, and lipid-associated metabolic regulation (58, 59). In parallel, reduced expression of transport-related genes, including *Slc7a2*, *Slc16a9*, *Cd320*, *Kcnj13*, and *Clcn5*, was consistent with pathway-level changes in solute transport, amino acid metabolism, and ionic homeostasis (**Figure 1**). These genes are associated with amino acid, monocarboxylate, vitamin B12, and ion transport, supporting the broader negative enrichment of transmembrane transport and amino acid metabolism pathways observed in exposed placentae **(Figure 3)**. In addition, negative enrichment of mTOR-related pathways may indicate altered regulation of nutrient-sensing programs, given the role of mTOR signaling in placental amino acid (60, 61). Together with our previous observation of reduced *Glut1* expression following prenatal cannabis smoke exposure, these findings support altered transcriptional regulation of placental transport-associated pathways (62). Furthermore, the placenta functions as a selective barrier that regulates fetal exposure to environmental insults (63). In response to xenobiotics, adaptive changes in placental transporter expression, including solute carrier (SLC)-mediated systems, may limit fetal exposure to potentially harmful compounds, such as constituents of cannabis smoke, potentially at the expense of nutrient availability (64). Taken together, these findings support a model in which prenatal cannabis smoke exposure induces coordinated changes in placental metabolic, bioenergetic, transport-associated and RNA processing gene programs.

A major strength of this study is the validation of selected microarray-identified differentially expressed genes using real-time qPCR in an expanded cohort that included the microarray samples, supporting the directionality of our transcriptomic findings (**Figure 4**). A further strength is the integration of pathway-level transcriptomic analyses with targeted mitochondrial follow-up experiments, including assessment of mitochondrial respiratory-chain gene expression, complex-associated respiration, and associated ETC complex protein content (**Figure 5**). An additional strength is the use of an inhalation-based exposure paradigm that more closely models real-world cannabis use during pregnancy and captures the integrated biological effects of whole cannabis smoke. Targeted cannabinoid quantification in paired maternal and fetal liver samples, performed as previously described, confirmed maternal and fetal cannabinoid delivery (21). Route of exposure is a critical determinant of cannabinoid pharmacokinetics and downstream biological effects, as inhalation introduces a complex mixture of combustion-derived compounds (65). While this complexity enhances the physiological relevance of our findings, it also presents challenges in attributing observed effects to specific cannabinoids, combustion-derived toxicants, or their interaction. Therefore, the transcriptomic changes identified here should be interpreted as the integrated placental response to combusted cannabis smoke rather than as evidence of cannabinoid-specific molecular effects. Future studies incorporating cannabinoid-depleted cannabis and/or combustion controls will be required to resolve the relative contribution of cannabinoid pharmacology and smoke-derived toxicants.

Despite these strengths, several limitations should be considered. The modest sample size may have limited statistical power to detect subtle gene-level differences and interaction effects, including sex-specific responses. Although fetal sex was included as a covariate, the study was not adequately powered to robustly assess sex-by-exposure interactions. Given increasing evidence for sex-specific placental adaptations in response to environmental stressors, this remains an important area for future investigation (66). Consistent with the limited sample size, we identified a modest number of differentially expressed genes. Therefore, pathway enrichment analyses were performed using ranked gene lists, such that enrichment reflects coordinated directional shifts in gene expression rather than absolute expression levels or uniform changes across all genes within a pathway. Related pathway terms may share substantial overlap in member genes and should not be interpreted as fully independent biological discoveries.

Accordingly, we interpret the recurrent enrichment of lipid metabolic, bioenergetic, and transport-related terms across annotation resources as convergent evidence for broader transcriptional themes rather than distinct pathway effects. While this approach does not account for baseline placenta-specific expression patterns, the concordance between pathway-level enrichment and gene-level changes supports the interpretation that these biological processes are differentially regulated in response to cannabis smoke exposure. Additionally, the use of bulk transcriptomic profiling limits the resolution of cell type-specific effects. As the placenta is a highly heterogeneous tissue, it is possible that cell-specific transcriptional changes were masked in our analysis, which may also explain the discrepancy with our GSVA and prior histological findings (67). Accordingly, the absence of large-scale separation in PCA, correlation, and hierarchical clustering analyses should not be interpreted as excluding exposure-driven transcriptional reprogramming within specific placental compartments or cell populations.

Furthermore, the limited availability of well-defined murine placental single-cell reference datasets presents challenges in constructing robust gene signatures. Future studies using single-cell and single-nuclei RNA sequencing approaches will be critical to resolve cell-type-specific responses and better define how distinct placental cell populations respond to cannabis smoke exposure.

In summary, prenatal cannabis smoke exposure was associated with subtle yet coordinated alterations in placental gene expression, including changes in compartment-associated transcriptional signatures, metabolic and bioenergetic pathways, transport-associated genes, RNA-processing programs, and selected mitochondrial respiratory-chain transcripts.

These transcriptional changes occurred alongside preserved complex-associated placental respiratory capacity and oxidative phosphorylation protein content. Collectively, this work provides new insight into placental responses to whole cannabis smoke exposure and highlights the importance of considering coordinated pathway-level and mitochondrial gene-regulatory changes, even in the absence of widespread differential gene expression or respiratory impairment.

## Acknowledgements

We thank the CRLB-GMEL laboratory, Population Health Research Institute (PHRI) for performing the microarray.

## Author Contributions

C.M., T.P., E.R., S.R. conceptualization; C.M., T.P., D.X., M.S., E.R., S.R. data curation; C.M., T.P., D.T., M.L. formal analysis. T.P., S.R. funding acquisition; C.M., T.P., D.X., M.L., A.M., M.S., S.W., S.R., E.R. methodology and investigation; C.M., T.P., D.X., D.T., M.L., S.W., S.R., interpretation; C.M., J.F., S.R., writing – original draft preparation; C.M., T.P., D.X., J.F., M.L., J.P., S.W., E.R., S.R. writing – reviewing and editing. S.R., supervision and project administration. All authors approved the final version of the manuscript.

## Funding

This work was supported by the Michael G. DeGroote Centre for Medicinal Cannabis Research at McMaster University and the Natural Sciences and Engineering Research Council (NSERC) Discovery Grant program (RGPIN-2020-06739).

## Data Availability Statement

The raw and processed data are available in the Gene Expression Omnibus with accession number GSE337449.

## Code Availability

The scripts used in this study are publicly available on GitHub at: https://github.com/crmonaco/Prenatal-Cannabis-Smoke-Placental-Transcriptomics/.

## Conflicts of Interest

The authors declare no competing interests.

## Notes

https://www.ncbi.nlm.nih.gov/geo/query/acc.cgi?acc=GSE337449

### Competing Interest Statement

The authors have declared no competing interest.

## References

1. Volkow ND, Han B, Compton WM, McCance-Katz EF. Self-reported Medical and Nonmedical Cannabis Use Among Pregnant Women in the United States. JAMA. 2019;322(2):167–9.

2. Watts D, Lebel C, Chaput K, Giesbrecht GF, Dewsnap K, Baglot SL, et al. Evaluation of the Association Between Prenatal Cannabis Use and Risk of Developmental Delay. JAACAP Open. 2024;2(4):250–62.

3. Young-Wolff KC, Cortez CA, Nugent JR, Padon AA, Prochaska JJ, Adams SR, et al. Sociodemographic differences in modes of cannabis use among pregnant individuals in Northern California. Drug Alcohol Depend. 2025;267:112546.

4. Gerede A, Stavros S, Chatzakis C, Vavoulidis E, Papasozomenou P, Domali E, et al. Cannabis Use during Pregnancy: An Update. Medicina (Kaunas). 2024;60(10).

5. Wiley JL, Taylor SI, Marusich JA. Δ. Drug Alcohol Depend. 2021;225:108827.

6. Black T, Baccetto SL, Barnard IL, Finch E, McElroy DL, Austin-Scott FVL, et al. Characterization of cannabinoid plasma concentration, maternal health, and cytokine levels in a rat model of prenatal Cannabis smoke exposure. Sci Rep. 2023;13(1):21070.

7. Baglot SL, VanRyzin JW, Marquardt AE, Aukema RJ, Petrie GN, Hume C, et al. Maternal-fetal transmission of delta-9-tetrahydrocannabinol (THC) and its metabolites following inhalation and injection exposure during pregnancy in rats. J Neurosci Res. 2022;100(3):713–30.

8. Black T, Barnard IL, Baccetto SL, Greba Q, Orvold SN, Austin-Scott FVL, et al. Differential effects of gestational Cannabis smoke and phytocannabinoid injections on male and female rat offspring behavior. Prog Neuropsychopharmacol Biol Psychiatry. 2025;136:111241.

9. Burton GJ, Fowden AL. The placenta: a multifaceted, transient organ. Philos Trans R Soc Lond B Biol Sci. 2015;370(1663):20140066.

10. Cindrova-Davies T, Sferruzzi-Perri AN. Human placental development and function. Semin Cell Dev Biol. 2022;131:66–77.

11. Napso T, Yong HEJ, Lopez-Tello J, Sferruzzi-Perri AN. The Role of Placental Hormones in Mediating Maternal Adaptations to Support Pregnancy and Lactation. Front Physiol. 2018;9:1091.

12. Vuppaladhadiam L, Lager J, Fiehn O, Weiss S, Chesney M, Hasdemir B, et al. Human Placenta Buffers the Fetus from Adverse Effects of Perceived Maternal Stress. Cells. 2021;10(2).

13. Walker OS, Ragos R, Gurm H, Lapierre M, May LL, Raha S. Delta-9- tetrahydrocannabinol disrupts mitochondrial function and attenuates syncytialization in human placental BeWo cells. Physiol Rep. 2020;8(13):e14476.

14. Podinic T, Limoges L, Monaco C, MacAndrew A, Minhas M, Nederveen J, et al. Cannabidiol Disrupts Mitochondrial Respiration and Metabolism and Dysregulates Trophoblast Cell Differentiation. Cells. 2024;13(6).

15. Monaco C, Minhas M, Podinic T, Nederveen JP, Lucas AM, Tomy T, et al. Cannabis smoke extract disrupts trophoblast differentiation and causes mitochondrial dysfunction beyond the effects of Δ9-THC alone. Sci Rep. 2026;16(1):6253.

16. Costa MA, Fonseca BM, Marques F, Teixeira NA, Correia-da-Silva G. The psychoactive compound of Cannabis sativa, Δ(9)-tetrahydrocannabinol (THC) inhibits the human trophoblast cell turnover. Toxicology. 2015;334:94–103.

17. Walker OS, Gurm H, Sharma R, Verma N, May LL, Raha S. Delta-9- tetrahydrocannabinol inhibits invasion of HTR8/SVneo human extravillous trophoblast cells and negatively impacts mitochondrial function. Sci Rep. 2021;11(1):4029.

18. Natale BV, Gustin KN, Lee K, Holloway AC, Laviolette SR, Natale DRC, et al. Δ9- tetrahydrocannabinol exposure during rat pregnancy leads to symmetrical fetal growth restriction and labyrinth-specific vascular defects in the placenta. Sci Rep. 2020;10(1):544.

19. Roberts VHJ, Schabel MC, Boniface ER, D’Mello RJ, Morgan TK, Terrobias JJD, et al. Chronic prenatal delta-9-tetrahydrocannabinol exposure adversely impacts placental function and development in a rhesus macaque model. Sci Rep. 2022;12(1):20260.

20. Graves BM, Johnson TJ, Nishida RT, Dias RP, Savareear B, Harynuk JJ, et al. Comprehensive characterization of mainstream marijuana and tobacco smoke. Sci Rep. 2020;10(1):7160.

21. Podinic T, Sunil M, MacAndrew A, Monaco C, Lee G, Lockington C, et al. Prenatal cannabis smoke exposure alters placental development in a murine model of pregnancy. PLoS One. 2026;21(3):e0328123.

22. Baker BH, Freije S, MacDonald JW, Bammler TK, Benson C, Carroll KN, et al. Placental transcriptomic signatures of prenatal and preconceptional maternal stress. Mol Psychiatry. 2024;29(4):1179–91.

23. Rosenfeld CS. Transcriptomics and Other Omics Approaches to Investigate Effects of Xenobiotics on the Placenta. Front Cell Dev Biol. 2021;9:723656.

24. Fantauzzi MF, Cass SP, McGrath JJC, Thayaparan D, Wang P, Stampfli MR, et al. Development and validation of a mouse model of contemporary cannabis smoke exposure. ERJ Open Res. 2021;7(3).

25. Ritchie ME, Phipson B, Wu D, Hu Y, Law CW, Shi W, et al. limma powers differential expression analyses for RNA-sequencing and microarray studies. Nucleic Acids Res. 2015;43(7):e47.

26. Irizarry RA, Hobbs B, Collin F, Beazer-Barclay YD, Antonellis KJ, Scherf U, et al. Exploration, normalization, and summaries of high density oligonucleotide array probe level data. Biostatistics. 2003;4(2):249–64.

27. Team RC. R: A language and environment for statistical computing. Vienna, Austria: R Foundation for Statistical Computing; 2024.

28. Phipson B, Lee S, Majewski IJ, Alexander WS, Smyth GK. ROBUST HYPERPARAMETER ESTIMATION PROTECTS AGAINST HYPERVARIABLE GENES AND IMPROVES POWER TO DETECT DIFFERENTIAL EXPRESSION. Ann Appl Stat. 2016;10(2):946–63.

29. Kolde R. pheatmap: Pretty Heatmaps. 2025.

30. Hänzelmann S, Castelo R, Guinney J. GSVA: gene set variation analysis for microarray and RNA-seq data. BMC Bioinformatics. 2013;14:7.

31. Simmons DG, Rawn S, Davies A, Hughes M, Cross JC. Spatial and temporal expression of the 23 murine Prolactin/Placental Lactogen-related genes is not associated with their position in the locus. BMC Genomics. 2008;9:352.

32. Jiang X, Wang Y, Xiao Z, Yan L, Guo S, Wu H, et al. A differentiation roadmap of murine placentation at single-cell resolution. Cell Discov. 2023;9(1):30.

33. Marsh B, Blelloch R. Single nuclei RNA-seq of mouse placental labyrinth development. Elife. 2020;9.

34. Van Buren E, Azzara D, Rangel-Moreno J, Garcia-Hernandez ML, Murphy SP, Cohen ED, et al. Single-cell RNA sequencing reveals placental response under environmental stress. Nat Commun. 2024;15(1):6549.

35. Wu T, Hu E, Xu S, Chen M, Guo P, Dai Z, et al. clusterProfiler 4.0: A universal enrichment tool for interpreting omics data. Innovation (Camb). 2021;2(3):100141.

36. Yu G, Wang LG, Han Y, He QY. clusterProfiler: an R package for comparing biological themes among gene clusters. OMICS. 2012;16(5):284–7.

37. Zyla J, Marczyk M, Weiner J, Polanska J. Ranking metrics in gene set enrichment analysis: do they matter? BMC Bioinformatics. 2017;18(1):256.

38. Ashburner M, Ball CA, Blake JA, Botstein D, Butler H, Cherry JM, et al. Gene ontology: tool for the unification of biology. The Gene Ontology Consortium. Nat Genet. 2000;25(1):25–9.

39. Kanehisa M, Furumichi M, Tanabe M, Sato Y, Morishima K. KEGG: new perspectives on genomes, pathways, diseases and drugs. Nucleic Acids Res. 2017;45(D1):D353–D61.

40. Liberzon A, Birger C, Thorvaldsdóttir H, Ghandi M, Mesirov JP, Tamayo P. The Molecular Signatures Database (MSigDB) hallmark gene set collection. Cell Syst. 2015;1(6):417–25.

41. Fabregat A, Jupe S, Matthews L, Sidiropoulos K, Gillespie M, Garapati P, et al. The Reactome Pathway Knowledgebase. Nucleic Acids Res. 2018;46(D1):D649–D55.

42. Livak KJ, Schmittgen TD. Analysis of relative gene expression data using real-time quantitative PCR and the 2(-Delta Delta C(T)) Method. Methods. 2001;25(4):402–8.

43. Podinic T, Xhuti D, Monaco C, Nederveen JP, Raha S. Assessing Mitochondrial Respiratory Complex-Associated Function From Previously Frozen Mouse Placental Tissue. Bio-protocol. 2026;16(9):e5667.

44. Delker E, Hayes S, Kelly AE, Jones KL, Chambers C, Bandoli G. Prenatal Exposure to Cannabis and Risk of Major Structural Birth Defects: A Systematic Review and Meta-analysis. Obstet Gynecol. 2023;142(2):269–83.

45. Woods L, Perez-Garcia V, Hemberger M. Regulation of Placental Development and Its Impact on Fetal Growth-New Insights From Mouse Models. Front Endocrinol (Lausanne). 2018;9:570.

46. Morasso MI, Grinberg A, Robinson G, Sargent TD, Mahon KA. Placental failure in mice lacking the homeobox gene Dlx3. Proc Natl Acad Sci U S A. 1999;96(1):162–7.

47. Clark PA, Brown JL, Li S, Woods AK, Han L, Sones JL, et al. Distal-less 3 haploinsufficiency results in elevated placental oxidative stress and altered fetal growth kinetics in the mouse. Placenta. 2012;33(10):830–8.

48. Chui A, Evseenko DA, Brennecke SP, Keelan JA, Kalionis B, Murthi P. Homeobox gene Distal-less 3 (DLX3) is a regulator of villous cytotrophoblast differentiation. Placenta. 2011;32(10):745–51.

49. Tan JL, Fogley RD, Flynn RA, Ablain J, Yang S, Saint-André V, et al. Stress from Nucleotide Depletion Activates the Transcriptional Regulator HEXIM1 to Suppress Melanoma. Mol Cell. 2016;62(1):34–46.

50. Li W, Wang L, Tian W, Ji W, Bing D, Wang Y, et al. SNRNP70 regulates the splicing of CD55 to promote osteosarcoma progression. JCI Insight. 2024;9(24).

51. Alves P, Amaral C, Gonçalves MS, Teixeira N, Correia-da-Silva G. Cannabidivarin and cannabigerol induce unfolded protein response and angiogenesis dysregulation in placental trophoblast HTR-8/SVneo cells. Arch Toxicol. 2024;98(9):2971–84.

52. Lojpur T, Easton Z, Raez-Villanueva S, Laviolette S, Holloway AC, Hardy DB. Δ9- Tetrahydrocannabinol leads to endoplasmic reticulum stress and mitochondrial dysfunction in human BeWo trophoblasts. Reprod Toxicol. 2019;87:21–31.

53. Kawahara T, Yanagi H, Yura T, Mori K. Endoplasmic reticulum stress-induced mRNA splicing permits synthesis of transcription factor Hac1p/Ern4p that activates the unfolded protein response. Mol Biol Cell. 1997;8(10):1845–62.

54. Toledano JM, Puche-Juarez M, Carrillo MP, Diaz-Castro J, Sanchez-Romero J, Ocaña- Peinado FM, et al. Placental dysregulation of mitochondrial morphology and dynamics as a hallmark of maternal age-related adaptation. Life Sci. 2026;393:124338.

55. Fisher JJ, McKeating DR, Cuffe JS, Bianco-Miotto T, Holland OJ, Perkins AV. Proteomic Analysis of Placental Mitochondria Following Trophoblast Differentiation. Front Physiol. 2019;10:1536.

56. Salazar-Petres E, Pereira-Carvalho D, Lopez-Tello J, Sferruzzi-Perri AN. Placental structure, function, and mitochondrial phenotype relate to fetal size in each fetal sex in mice†. Biol Reprod. 2022;106(6):1292–311.

57. Campbell KA, Colacino JA, Puttabyatappa M, Dou JF, Elkin ER, Hammoud SS, et al. Placental cell type deconvolution reveals that cell proportions drive preeclampsia gene expression differences. Commun Biol. 2023;6(1):264.

58. Shah DI, Takahashi-Makise N, Cooney JD, Li L, Schultz IJ, Pierce EL, et al. Mitochondrial Atpif1 regulates haem synthesis in developing erythroblasts. Nature. 2012;491(7425):608–12.

59. Shetty T, Sishtla K, Park B, Repass MJ, Corson TW. Heme Synthesis Inhibition Blocks Angiogenesis via Mitochondrial Dysfunction. iScience. 2020;23(8):101391.

60. Hart B, Morgan E, Alejandro EU. Nutrient sensor signaling pathways and cellular stress in fetal growth restriction. J Mol Endocrinol. 2019;62(2):R155–R65.

61. Beetch M, Oribamise E, Jo S, Clifton B, Larson S, Hausmann A, et al. Placental mTOR signalling links mitochondrial dysfunction, nutrient transport and neonatal beta cell perturbations in mice. Diabetologia. 2025;68(12):2823–39.

62. Jentsch TJ. Chloride and the endosomal-lysosomal pathway: emerging roles of CLC chloride transporters. J Physiol. 2007;578(Pt 3):633–40.

63. Mathiesen L, Buerki-Thurnherr T, Pastuschek J, Aengenheister L, Knudsen LE. Fetal exposure to environmental chemicals; insights from placental perfusion studies. Placenta. 2021;106:58–66.

64. Walker N, Filis P, Soffientini U, Bellingham M, O’Shaughnessy PJ, Fowler PA. Placental transporter localization and expression in the Human: the importance of species, sex, and gestational age differences†. Biol Reprod. 2017;96(4):733–42.

65. Chayasirisobhon S. Mechanisms of Action and Pharmacokinetics of Cannabis. Perm J. 2020;25:1–3.

66. Cowell W, Deyssenroth M, Chen J, Wright RJ. Maternal stress in relation to sex-specific expression of placental genes involved in nutrient transport, oxygen tension, immune response, and the glucocorticoid barrier. Placenta. 2020;96:19–26.

67. Zhou X, Xu Y, Ren S, Liu D, Yang N, Han Q, et al. Single-cell RNA-seq revealed diverse cell types in the mouse placenta at mid-gestation. Exp Cell Res. 2021;405(2):112715.

