## Supplemental Fig S1 and Table 1-2 for "Placental transcriptomics reveals alterations in metabolic and transport gene programs following prenatal cannabis smoke exposure"

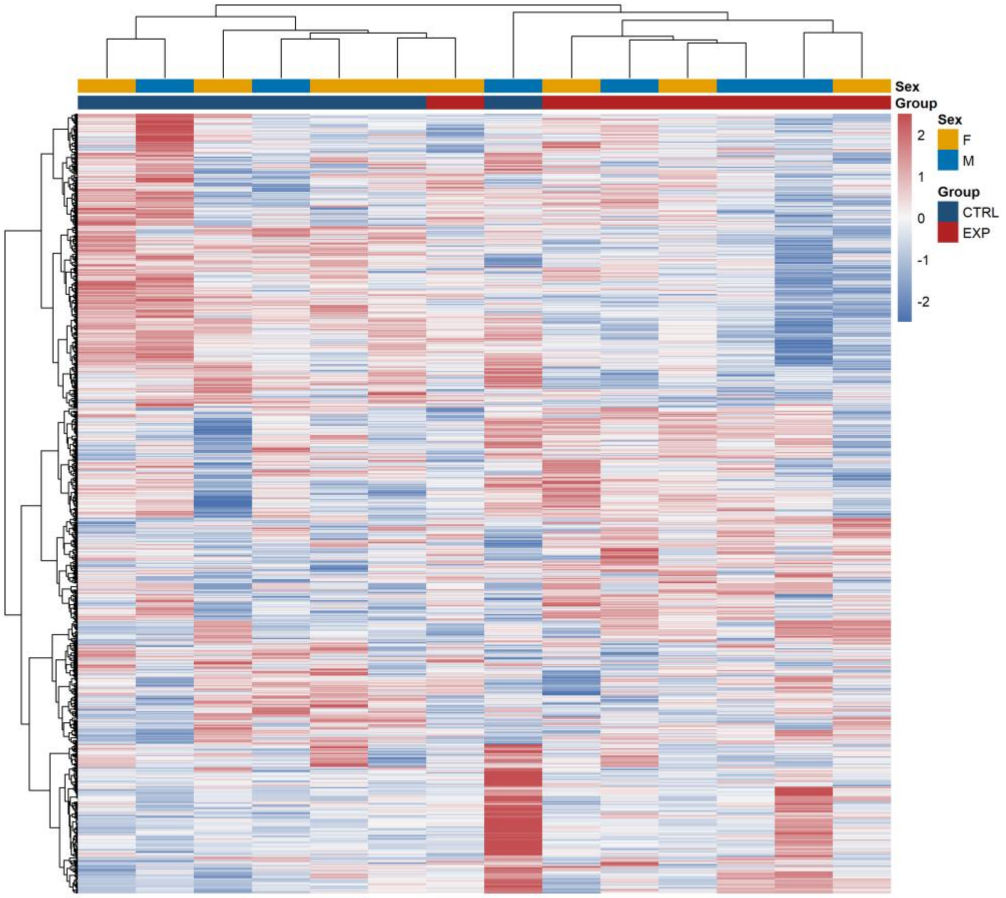


**Figure S1. Unsupervised hierarchical clustering of highly variable placental transcripts.** Heatmap displaying the top 1,000 most variable genes across E18.5 placental samples from control (CTRL) and cannabis smoke-exposed (EXP) groups. Gene expression values were row-scaled by z-score transformation prior to clustering. Columns represent individual placental samples and rows represent genes. Hierarchical clustering was performed using Euclidean distance and Ward's D2 linkage. Sample annotations indicate fetal sex (F, female; M, male) and exposure group (CTRL, control; EXP, cannabis smoke-exposed).

**Supplementary Table S1: Primer sets used in endpoint and real-time qPCR**

| Primer Set | Forward Primer | Reverse Primer |
|---|---|---|
| Sry | TTGTCTAGAGAGCATGGAGGGCCATGTCAA | CCACTCCTCTGTGACACTTTAGCCCTCCGA |
| Fabp2 | TGGACAGGACTGGACCTCTGCTTTCCTAGA | TAGAGCTTTGCCACATCACAGGTCATTCAG |
| ActB | AGCCATGTACGTAGCCATCCA | TCTCCGGAGTCCATCACAATG |
| Clcn5 | GGAAGAAACAGGACGGGGTT | GGTCTGTCACAGTGAAGGGG |
| Kcnj13 | ACGAGCCAGCACAATCATCT | GTGACCTGCTGCTGTCTGTA |
| Gypa | AACCACTCAGAGTACCCCCT | TAACGCTTTCACCCATGCCT |
| Slc7a2 | CTGTTCCTTGTCCTCTGCGT | TTCTGTGGCTGCCTCCAAAT |
| Slc16a9 | TACATTGCCTTCCTTTGGGGA | GGTGCTATCCGGGATGTGTAA |
| Tcf20 | TGGAGACCACTGCCATCCTA | TGCATTGCCACGGTAAGTCT |
| Ndufb8 | ATGTTGCCGGGGTCATATCC | GGCTCGTAGTCTTCCACTCG |
| Ndufs1 | TCTTGCTGACCCACTCGTTC | CGGTGACAGCTTTGACACAC |
| Ndufs2 | AGCGAGCAGAGATGAAGACG | CTGGAGGAACTTGGTAGCCC |
| Ndufv1 | CCACACCCTAGCCTGACAAT | GTTGGAGAGTGGACAGCACT |
| Sdha | GCAGTTTCGAGGCTTCTTCG | AAGCCGCAGGTCTGTTTTTG |
| Sdhb | GGACCTATGGTGTTGGATGCT | GCCTCCGTTGATGTTCATGG |
| Uqcrc1 | AAGTTAGAAGATGGCGGCGT | CAAGGCAGGTAACCTCAGCA |
| Uqcrc2 | AGGTTAAAACTTCAGCAGCCC | ACGAACAAGCCGATTCTTGAC |
| Cox4i1 | TGACTACCCCTTGCCTGATG | ACTGGATGCGGTACAACTGA |
| Cox5a | GCTGTCTGTTCCATTCGCTG | CCTTTACGCAATTCCCAGGC |
| Cox7a2 | CCTTCGTCAGATTGCCCAGA | AATGCCTTCGTGAAGTGGTG |
| Atp5f1a | AGAATCGCCTGGACTACCAC | GCAATGGTCTCCTTTTCCTGC |
| Atp5o | CTTGCTGAAAATGGTCGCCT | GAGGAGATGCTGTGGTCACT |
| Pparag | CCCTGCCATTGTTAAGACC | TGCTGCTGTTCCTGTTTTC |
| Tfam | ATGTGGAGCGTGCTAAAAGC | TGGGTAGCTGTTCTGTGGAAA |

### Supplementary Table S2: Top statistically significant genes (FDR p<0.05)

| Symbol | Gene Name | log2 FC | FDR p-value |
|---|---|---|---|
| Tcf20 | transcription factor 20 | 1.24 | 0.0271 |
| 6030498E09Rik | RIKEN cDNA 6030498E09 gene | 0.98 | 0.0271 |
| Hexim1 | hexamethylene bis-acetamide inducible 1 | 0.95 | 0.0309 |
| Snrnp70 | small nuclear ribonucleoprotein 70 (U1) | 0.91 | 0.0413 |
| Sumo3 | small ubiquitin-like modifier 3 | 0.9 | 0.0324 |
| Kcnj13 | potassium inwardly rectifying channel, subfamily J, member 13 | -2.11 | 0.00297 |
| Gypa | glycophorin A | -1.94 | 0.0271 |
| Slc7a2 | solute carrier family 7 (cationic amino acid transporter, y+ system), member 2 | -1.74 | 0.00297 |
| Cd320 | CD320 antigen | -1.72 | 0.0271 |
| Fech | ferrochelatase | -1.62 | 0.0358 |
| Slc16a9 | solute carrier family 16 (monocarboxylic acid transporters), member 9 | -1.58 | 0.0271 |
| Tspan1 | tetraspanin 1 | -1.52 | 0.0271 |
| Crb3 | crumbs family member 3 | -1.47 | 0.0271 |
| Clcn5 | chloride channel, voltage-sensitive 5 | -1.35 | 0.0318 |
| Lpcat3 | lysphosphatidylcholine acyltransferase 3 | -1.29 | 0.047 |
| Prcp | prolylcarboxypeptidase (angiotensinase C) | -1.21 | 0.0271 |
| Samhd1 | SAM domain and HD domain, 1 | -1.19 | 0.0271 |
| Ccdc115 | coiled-coil domain containing 115 | -1.12 | 0.0286 |
| Dlx3 | distal-less homeobox 3 | -1.09 | 0.0271 |
| Got1 | glutamic-oxaloacetic transaminase 1, soluble | -1.07 | 0.0354 |
| Dvl3 | dishevelled segment polarity protein 3 | -1.03 | 0.0271 |
| Elovl1 | ELOVL fatty acid elongase 1 | -1.02 | 0.0354 |
| Nit1 | nitrilase 1 | -1.01 | 0.0271 |

*FC, fold change; FDR; false discovery rate

**Supplementary Table S3. Curated cluster-resolved trophoblast markers for labyrinth and junctional zone markers used in GSVA**

| Genes | Function | Compartment | Source |
|---|---|---|---|
| Tpbpa | Central progenitor marker | Junctional | Jiang et al., 2023 (37); Van Buren et.al., 2024 (39); Marsh & Blelloch, 2020 (38); Simmons et al., 2008 (36) |
| Prl3d1, Prl2c2, Prl3b1 | Terminal endocrine trophoblast marker | Junctional | Jiang et al., 2023 (37); Van Buren et.al., 2024 (39); Marsh & Blelloch, 2020 (38); Simmons et al., 2008 (36) |
| Prl7b1, Prl8a8 | Prolactin-family markers | Junctional | Jiang et al., 2023 (37); Van Buren et.al., 2024 (39); Marsh & Blelloch, 2020 (38); Simmons et al., 2008 (36) |
| Pcdh12 | Glycogen trophoblast marker | Junctional | Jiang et al., 2023 (37); Van Buren et.al., 2024 (39); Marsh & Blelloch, 2020 (38); Simmons et al., 2008 (36) |
| Gcm1 | Master regulator of syncytiotrophoblast | Labyrinth | Jiang et al., 2023 (37); Marsh & Blelloch, 2020 (38) |
| Syna, Synb | Fusion genes for SynT | Labyrinth | Jiang et al., 2023 (37) |
| Epcam | Labyrinth progenitor (LaTP) | Labyrinth | Jiang et al., 2023 (37); Van Buren et.al., 2024 (39); Marsh & Blelloch, 2020 |
| Dlx3 | Placental differentiation | Labyrinth | Jiang et al., 2023 (37); Marsh & Blelloch, 2020 (38) |
| Slc16a3 (Mct4) | Nutrient/lactate transport | Labyrinth | Jiang et al., 2023 (37) |
| Msx2 | Labyrinth differentiation transcription factor | Labyrinth | Jiang et al., 2023 (37); Hornbachner et al., 2021 (73) |
| Ovol2 | Trophoblast differentiation | Labyrinth | Jiang et al., 2023 (37); Jeyarajah et al., 2020 (74) |
